# Siglec homologs interact with membrane-bound Heat Shock Protein 70 (Hsp70) during early infection of the *Schistosoma mansoni* susceptible (BB02) *Biomphalaria glabrata snail* host

**DOI:** 10.1101/2025.07.18.665599

**Authors:** Eli Vanlal, Oumsalama Elhelu, Olayemi G. Fagunloye, Matty Knight

## Abstract

Sialic acid-binding immunoglobulin-like lectins (siglecs) are cell surface receptors involved in immune signaling. When *Schistosoma mansoni* infects its intermediate host, the *Biomphalaria glabrata* snail, early stress responses, such as upregulation of heat shock protein 70 (Hsp70) are observed. We hypothesized that stress-induced Hsp70 translocates to the cell membrane and interacts with siglec homologs to modulate immune responses in the snail. Utilizing in-silico approaches, we identified *B. glabrata* transcripts homologous to human and molluscan siglecs, followed by homology modeling and molecular docking, which predicted a salt bridge between glutamic acid 479 on *BgHsp70* and lysine 47 on a siglec homolog (*BgPrx*), suggesting a plausible binding interface. To validate this, we performed real-time qPCR from infected susceptible juvenile BB02 snails, revealing significant upregulation of siglec homolog transcripts shortly after infection. Additionally, protein fractionation and immunocytochemistry confirmed *BgHsp70* localization to the membrane post-infection. These results support a model in which siglec–Hsp70 interactions may dampen stress signaling to suppress host immune defenses. Concurrently, *S. mansoni* employs glycan mimicry, presenting host-like sialylated structures that likely engage siglecs and further misdirect the immune response.

Together, our findings suggest that *BgHsp70*–siglec interactions, in combination with parasite glycan mimicry, constitute a potential immune evasion mechanism enabling schistosome establishment in susceptible *B. glabrata* snails.

## INTRODUCTION

The freshwater pulmonate gastropod, *Biomphalaria glabrata*, is the obligate intermediate host for the development of the parasitic trematode, *Schistosoma mansoni*, in the Western Hemisphere. The parasite is the etiological agent for the chronic debilitating tropical disease schistosomiasis, which is commonly known as either bilharzia or snail fever. Schistosomiasis is prevalent across the developing countries of sub-Saharan Africa, the Americas, the Arabian Peninsula, the Mediterranean, and the Pacific Rim. It is a disease of poverty in regions of the world where access to clean water and sanitation is lacking. It is estimated that at least 600 million people globally are at risk of infection if they live in endemic regions close to freshwater bodies such as lakes, rivers, and their tributaries [1]. Schistosomiasis is known as the second most significant debilitating tropical disease in the world, after malaria [2]. Without a vaccine, prevention of schistosomiasis remains elusive, and the single drug, praziquantel, that is used to treat the disease, is only efficacious against adult parasites, not their larval forms. Therefore, without new drugs or vaccines to control the spread of the disease in endemic regions, eliminating schistosomiasis remains a major challenge. Lately, in response to the World Health Organization’s call to reduce prevalence and eliminate schistosomiasis by 2025, efforts to control disease transmission have shifted towards exploring new approaches to interfere with the development of the parasite in the intermediate snail host [3].

While the life cycle of the parasite is complex, requiring both human and intermediate snail hosts for its development (Figure 1), progress is being made to better understand and identify the key determinants that trigger an anti-schistosome response in the snail host [4]. Previously, we showed that an early stress response mounted in juvenile snails to invading schistosomes is a hallmark of its susceptibility to the parasite [5]. The relationship between the snail and parasite has been shown to be highly variable with several factors, including age, affecting *B. glabrata* susceptibility to *S. mansoni*. Thus, the same snail, susceptible as a juvenile can become resistant as it ages, and a resistant snail can also become susceptible if bred at an elevated temperature of 32°C [6-8]. These data showing intrastrain variability in the *B. glabrata-S. mansoni* relationship with either age or elevated temperature remain intriguing. For example, we recently showed that epigenetics involving an interplay between PIWI and the endogenous *B. glabrata* non-LTR retrotransposon, *nimbus*, plays a yet unknown mechanism in resistance to schistosome infection in the snail host [8]. Furthermore, an innate defence system involving a variety of immunodominant plasma lectins that act in the self/non-self-recognition together with hemocytes to engulf and destroy the invading schistosome in a non-compatible snail host has long been documented by various investigators [9, 10], [11].

**Figure 1.**
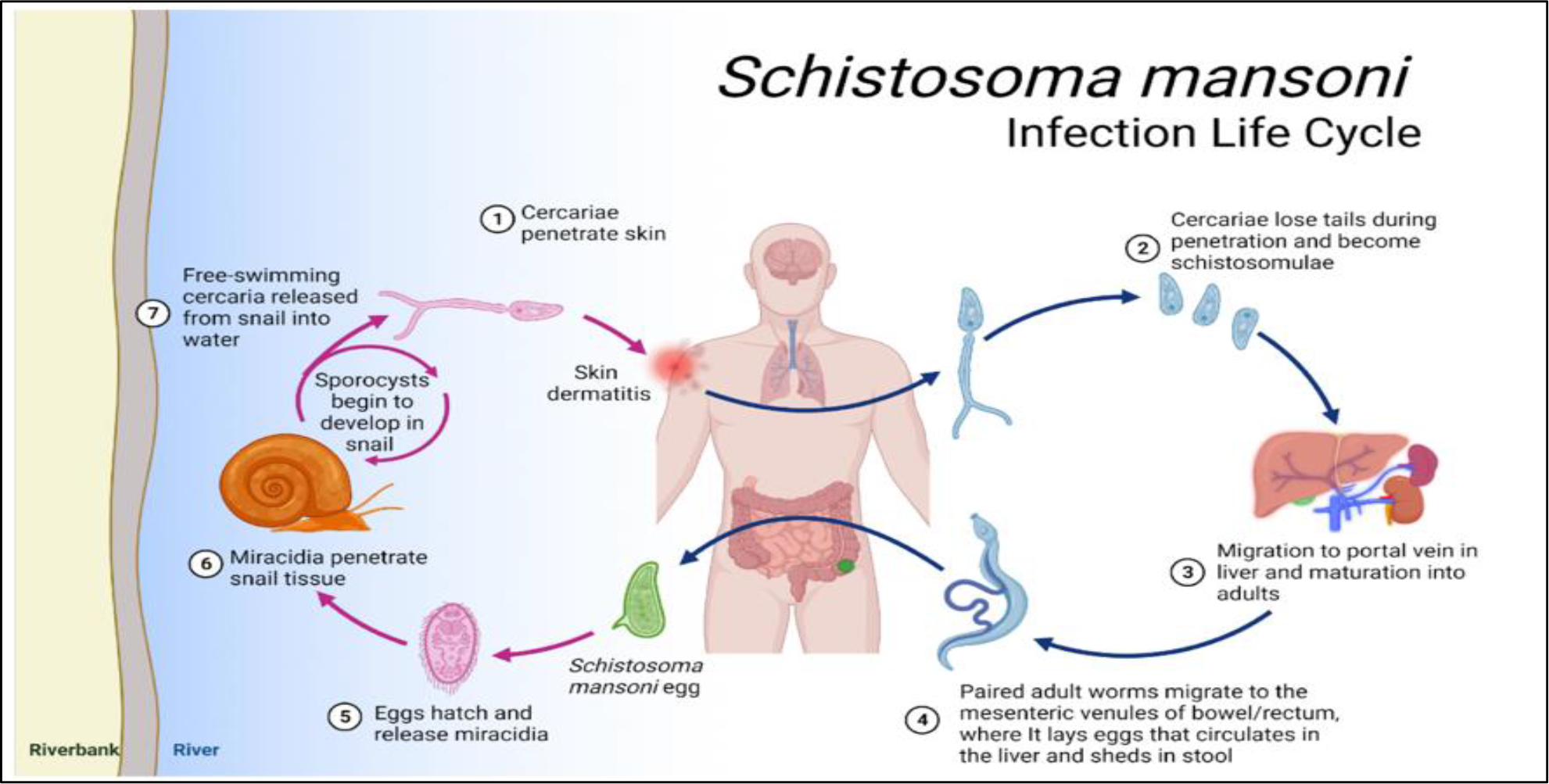
Life cycle of *Schistosoma mansoni* highlighting developmental stages in the human and intermediate snail host, *Biomphalaria glabrata*. Cercariae swimming in infested water body penetrate human skin and become schistosomula **(1–2)** then migrate to the liver and mature into adult worms **(3)**. Adults reside in mesenteric veins, releasing eggs that exit via stool (4–5). In freshwater, eggs hatch into miracidia that infect snails **(6)**, where sporocyst development occurs. Cercariae are later released from the snail to continue the cycle **(7)**.

The past decade has revealed a plethora of factors that play an important role in the complex response of the *B. glabrata* snail-schistosome interaction; an ancient relationship suggested to be the driving force, rather than the vertebrate host, in shaping the genetics of evolutionary behavior/adaptation of parasitic trematodes in both snail and human hosts [12, 13] [14-16]. Thus, in this era of omics-based research, the availability of a reference genome sequence of *B. glabrata* [17], along with transcriptomics and proteomics of various snail tissues (including single-cell hemocytes), provides an opportunity to identify important snail determinants that interact with miracidia to either protect or destroy the developing parasite. Here, we reveal how without any prior sequence data, siglec homologs of *B. glabrata* have been identified entirely by utilizing an omics approach. We also show that these *B. glabrata* siglec homologs are induced in the early *S. mansoni* response in juvenile susceptible *B. glabrata* snails and for the first time, report the induction of membrane-bound Hsp70 and its interaction with the siglec homolog of *B. glabrata* peroxidasin.

Siglecs, attach to glycans on cell surfaces. They are important in immune cell signaling, acting as either positive or negative regulators of the innate immune system. When the miracidia of *Schistosoma mansoni* infect the freshwater snail, *Biomphalaria glabrata*, stress proteins are elevated. This deploys an immune response that upregulates the expression of variable immunoglobulin-like lectins, such as siglec homologs. The developing miracidia evade lysis by disguising themselves as glycan structures like those found on host immune cells. These glycans can bind to siglecs, providing effective mimicry that prevents recognition by the snail’s innate defence system.

This evasive mechanism, utilized by larval schistosomes to escape recognition and destruction from the snail’s innate defence system to enable the invading miracidiae to survive in what should be a hostile environment, remains poorly understood. Hsp70 has been described as a cell surface receptor that associates with siglecs to modulate the host immune response [18].

To determine the mechanism(s) that the snail host uses in this anti-schistosome innate defence response, we hypothesized that siglecs in *B. glabrata* are upregulated in susceptible snails. We also rationalized that a possible surface location of *B. glabrata* Hsp70 in association with siglecs would aid the cell-to-cell adhesion that is required for the hemocyte encapsulation of miracidia during the anti-schistosome response in the snail.

To test this hypothesis, we used an *in-silico* approach to identify *B. glabrata* siglec homologs by interrogating the snail’s genome with amino acid sequences corresponding to known human and mollusc siglecs in the amino acid database, UniProt, and also in the published literature, followed by a BLASTp analysis in NCBI. These searches confirmed the existence of putative *B. glabrata* siglecs showing a significant match with their human and molluscan counterparts. For example, the amino acid of human Siglec-15 shared a significant homology with *B. glabrata* peroxidasin. Gene-specific primers were designed from the corresponding *B. glabrata* sequences and utilized to examine the temporal regulation of expression by qPCR. Results showed that following early infection, transcripts encoding siglec homologs were upregulated in susceptible *B. glabrata (*BBO2 stock). A better understanding of how these putative siglec homologs provide either innate defense, camouflage, or immune evasion in the anti-schistosome response in *B. glabrata* will be an important step toward the development of novel snail vector-based transmission blocking strategies for eliminating schistosomiasis.

## MATERIALS AND METHODS

### Snails

Juvenile *B. glabrata* BB02 – susceptible snails provided by the NIAID schistosomiasis resource center (https://www.afbr-bri.org/contact-bri/) were utilized throughout the study. The snails (3.0 to 4.0cm in diameter) were exposed, individually, to *S*.*mansoni* (NMRI strain)10 miracidia per snail in 2.0ml of aerated tap water in six-well microtiter plates for specific time points (0, 30 mins, 1hr., 2 hr., 4 hr. and overnight). After infection, snails were either processed immediately for RNA isolation or kept at -80°C until required.

### RNA Isolation

Fresh or snails frozen in 500 ul of RNAzol at – 80°C were homogenized in microcentrifuge tubes as previously described [8].

### Bioinformatics

FASTA files corresponding to amino acid sequences of known human and molluscan siglecs were obtained either from the protein data base uniport (www.uniprot.org) or the published literature (https://pubmed.ncbi.nlm.nih.gov/). Sequences of interest were utilized as query to interrogate the *B. glabrata* genome in NCBI (BLAST https://blast.ncbi.nlm.nih.gov/Blast.cgi), while excluding *Schistosoma mansoni* to eliminate parasite related sequences. The known sialic acid binding lectins identified in molluscs’ such as *Haliotis discus discus;* (Accession no. AB026662.1) and *Cepaea hortensis* (Accession no. AJ551448) showed that C1q, peroxidasin, and nuclear receptor are significant *B. glabrata* siglec homologs. Sequences obtained from NCBI were used to design corresponding primers (Table 1) using primer BLAST in NCBI.

**Table 1.**
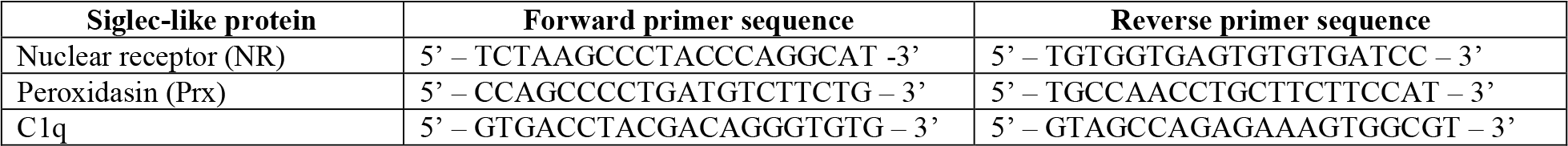
Primers for putative *Bg* siglec Gene Specific Sequences.

### cDNA Synthesis and Real time PCR

cDNA was synthesized from 500ng of each RNA sample using iScript cDNA synthesis kit and a thermal cycler PCR machine as previously described [8].

Real time PCR was performed by 2-step RT qPCR method to quantitatively assess the expression transcripts. We used SYBR Green PCR Master Mix purchased from Applied Bio systems (Thermo Fisher Scientific, Wolston Warrington, UK), and 15uM each of forward and reverse primers listed in Table 1 to evaluate the temporal expression of the *B. glabrata* putative siglec homologs with cDNA templates prepared from either resistant (BS-90) or susceptible (BBO2) snails.

Results were obtained from four biological replicates with (n=72) with each sample run in triplicate normalized against the myoglobin housekeeping gene which is constitutively expressed in the snail. A 7300 thermal cycler system was used to quantitatively measure the genes of interest using the relative quantification to evaluate the ΔΔCt. The resulting fold change in expression of the genes of interest were calculated by using the formula below [19]:

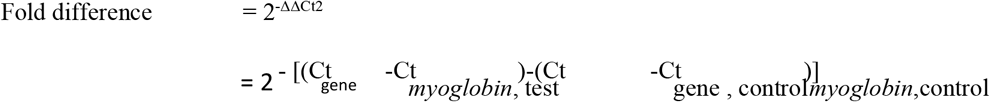

*P*-values were calculated by comparing the delta-Ct value for each sample and Student’s *t*-test was used to determine if there was significant difference in the differential expression of the transcripts between experimental and control groups with Graph Pad prism software (Version 9.4.1).

### Cytosolic and Membrane-Bound Protein Extraction

Juvenile snails were each exposed to 15 miracidia in six-well microtiter plates for varying durations (0 min, 30 min, 1 h, and 2 h). Following exposure, snails were homogenized in ice-cold 1x phosphate-buffered saline (PBS, pH 7.4) supplemented with protease inhibitor cocktail (Roche, Cat. No. 05056489001), then centrifuged at 13,000 rpm for 10 minutes at 4 °C. The supernatant, containing cytosolic proteins, were collected into fresh tubes. The remaining pellet was re-homogenized in radioimmunoprecipitation assay (RIPA) buffer (Sigma Millipore), supplemented with protease inhibitor cocktail (Roche, Cat. No. 05056489001), followed by a second centrifugation at 13,000 rpm for 10 minutes at 4 °C. The resulting supernatant, enriched in membrane-associated proteins, was collected separately.

### Western Blot

Soluble cytosolic and membrane-enriched protein extracts were prepared from exposed and unexposed snails at multiple time points post-infection. Ten micrograms of protein per sample were mixed with 4x Laemmli Sample Buffer (Bio-Rad, Cat. No. 1610747) supplemented with 10% β-mercaptoethanol (v/v). Samples were then dry-heated at 70°C for 10 minutes. Proteins were separated by SDS-PAGE on 4–10% gradient gels (Bio-Rad) using standard 1x Tris/Glycine/SDS running buffer. After electrophoresis, proteins were transferred onto nitrocellulose membranes using the Trans-Blot® Turbo™ Transfer System (Bio-Rad). Membranes were air-dried for 10 minutes at 37°C, stained with Revert™ 700 Total Protein stain (LI-COR), and then imaged using a fluorescence-based imaging system for normalization. Membranes were then blocked and incubated overnight at 4°C with rabbit -anti Hsp70 primary antibody (1:1000), followed by incubation with fluorophore-conjugated donkey anti-rabbit secondary antibody (LiCOR IRDye 800 CW). This Western blot analysis was used to qualitatively and quantitatively determine the occurrence of Hsp70 expression between cytosolic and membrane-associated protein fractions in exposed and unexposed snails.

### Preparation of *B. glabrata* Hsp 70 Antibody

Polyvalent rabbit antibody was made against the recombinant *B. glabrata* Hsp70 protein expressed in *E*.*coli* by Abclonal.

### Immunocytochemistry of exposed and unexposed *B. glabrata* embryonic cells (BGE)

*Bge* cells maintained in the laboratory were passaged and seeded onto 96-well optical-bottom plates (Greiner). Cells were allowed to adhere for 48 hours prior to exposure. For infection, cells were incubated with 10 - 15 *S. mansoni* miracidia for 2 hours. Following exposure, cells were fixed in 4% paraformaldehyde and permeabilized using 0.4% Triton X-100 in PBS. Cells were then incubated overnight at 4°C with anti-Hsp70 primary antibody (1:1000 dilution, rabbit polyclonal). After PBS washes, nuclei were stained using 10 µg/mL Hoechst 33342. Alexa Fluor 555-conjugated donkey anti-rabbit IgG (1:1000; Life Technologies, Ref. A31572) was used as the secondary antibody. Imaging was performed using an epifluorescence-based high-content microscope with confocal capabilities.

### Homology modeling and protein-protein docking

Homology models for *Biomphalaria glabrata* Hsp70 and a siglec homolog, proxidasin, were generated using SWISS-MODEL [31]. The FASTA sequence for BgHsp70 (accession: AAB95297.1) was submitted, and SWISS-MODEL selected the *Cristaria plicata* Hsp70 (UniProt accession: E1B2C9.1.A) as the modeling template, which shared 78.23% sequence identity. For the *B. glabrata* Siglec homolog (sequence XP_013091598.1) annotated as a peroxidasin-like protein, was submitted to SWISS-MODEL. The tool selected an AlphaFold-predicted structure of a *Cepaea hortensis* siglec homolog (Q70SH0.1.A) as the template, based on 100% sequence identity.

Protein–protein docking was performed using GRAMM [26], which generated six docking prediction models. Binding affinity (ΔG), dissociation constants (Kd) and interface contacts, were calculated to evaluate the strength and stability of the predicted interactions by submitting the files of the six models to PRODIGY [28, 29]. The docking interfaces were further visualized and analyzed using Mol* to identify interfacial residues and characterize potential interactions [30]

## RESULTS AND DISCUSSION

### *B. glabrata* peroxidasin is homologous to human siglec 15

To identify the putative homologs of siglecs in *B. glabrata*, amino acid sequences corresponding to known human siglecs, such as Human siglec 15 (alternative name: CD33 antigen-like 3) identified in UniProt (www.uniprot.org) was utilized.

In searching the Basic Local Alignment Search Tool (BLASTp) in the National Center for Biotechnology Information (NCBI-GenBank) with *B. glabrata* genomic sequences as query in this public data base, results revealed significant homology between the amino acid of human siglec 15 and *B. glabrata* peroxidasin (Accession no. XP_013091598.1). Results from BLASTp analysis, showed the amino acid sequence of the *B. glabrata* homolog was 30.19% identical to human siglec15 over 43% of the protein sequence with an E value of 5e^-5^

### *B. glabrata* C1q amino acid sequences are homologous to SABL in other molluscs

Similar interrogation of the *B. glabrata* genome with the amino acid sequence (Accession number: CAD83837.1) of the mollusc, *Cepaea hortensis* revealed a significant match (E-value 6e^-17^) of this sequence to *B. glabrata* complement C1q like protein 3 (XP_013081758.1) and protein 4 (XP_013091954.1, E value 3e^-12^). A similar BLASTp search, using amino acid sequence of the sialic acid binding lectin (SABL) of *Haliotis discus discus* (Fig. 3B.; Assession number ABO26662.1) also showed that this sequence is homologous to *B. glabrata* C1q like protein 3 (XP_013081758.1, E value 2e^-38^), protein 4 (XP_013091313.2, E value 1e^-22^), and C1q tumor necrosis factor-related protein 7 (KA18775437.1 E value 6e^-18^), annotated as nuclear receptor (NR), in Vector Base (http://vectorbase.org). Results of either human or molluscan amino acid sequences of siglecs that were found to match *B. glabrata* proteins in GenBank are listed in Table 2.

**Table 2.** siglecs with significant homology to *B. glabrata*.

| Description | Max score | Total score | Query cover | E Value | Identity | Accession |
| --- | --- | --- | --- | --- | --- | --- |
| Sialic acid-binding Ig-like lectin 15 X1<br>[*Mus musculus*] | 491 | 491 | 96% | 9e-175 | 83.65% | XP_030106421.1 |
| V-set and Ig domain-containing protein precursor<br>[*Danio rerio*] | 157 | 157 | 64% | 1e-44 | 38.69% | NP_001095865.1 |
| Sialic acid binding Ig-like lectin 7 isoform X1<br>[*Homo sapiens*] | 65.9 | 65.9 | 66% | 1e-10 | 25.11% | XP_011525023.1 |
| Leucine-rich repeats and immunoglobulin-like domains protein<br>sma-10 [Caenorhabditis elegans] | 53.1 | 53.1 | 79% | 3e-06 | 26.52% | NP_001367941.1 |
| Sialic acid-binding Ig-like lectin 15 precursor<br>[Mus musculus] | 486 | 486 | 96% | 6e-173 | 83.39% | NP_001094508.1 |

**Figure 2.**
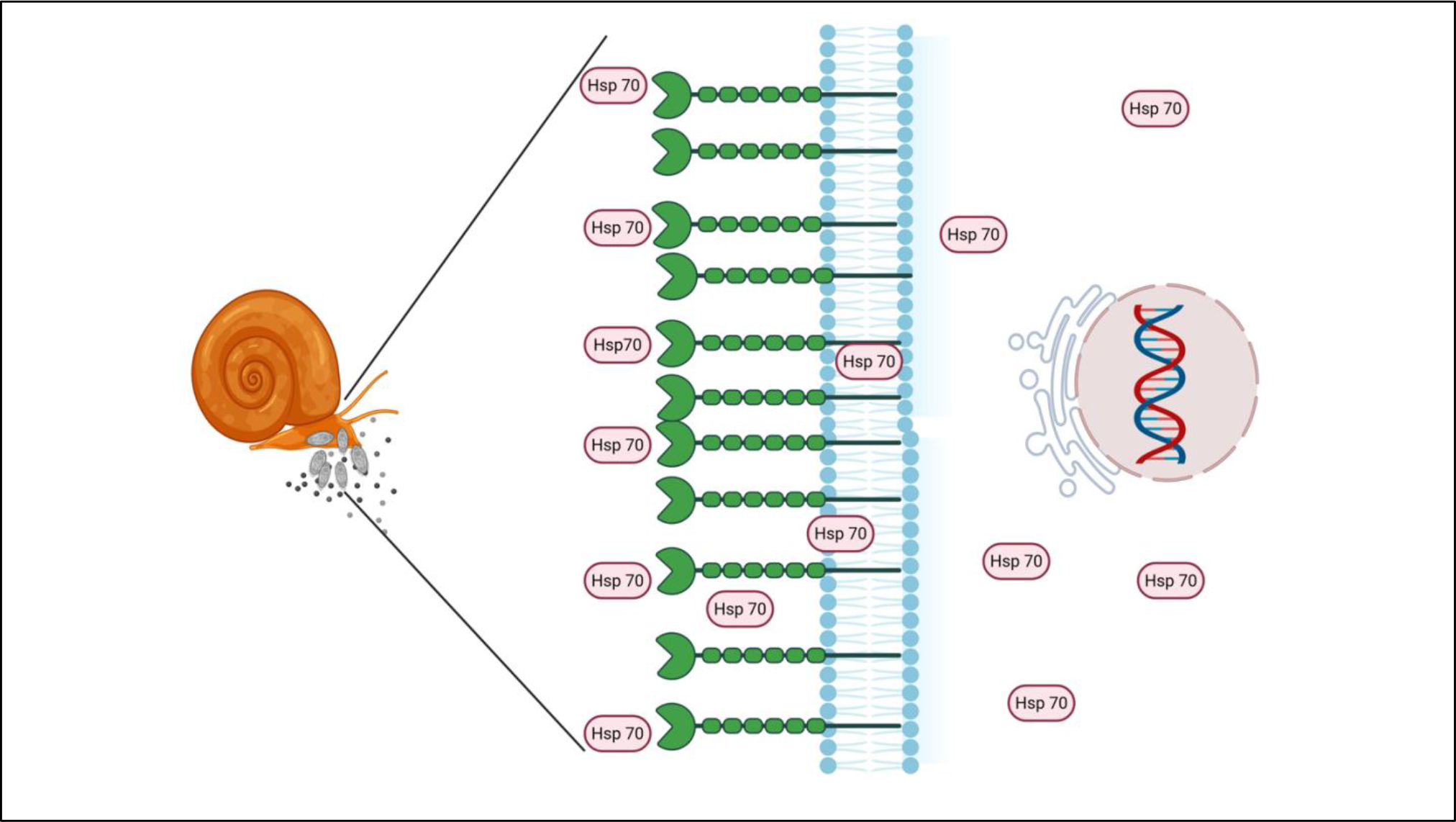
Proposed model of Hsp70 interaction with siglec homologs on the hemocyte membrane of *Biomphalaria glabrata*. Upon exposure to *Schistosoma mansoni* miracidia, Hsp70 translocates to the hemocyte surface, where it associates with putative siglec receptors (green) embedded in the membrane. This interaction may promote hemocyte cell-cell adhesion and encapsulation of invading miracidia as part of the snail’s innate immune response. Intracellular Hsp70 also contributes to transcriptional regulation and stress signaling.

**Figure 3.**
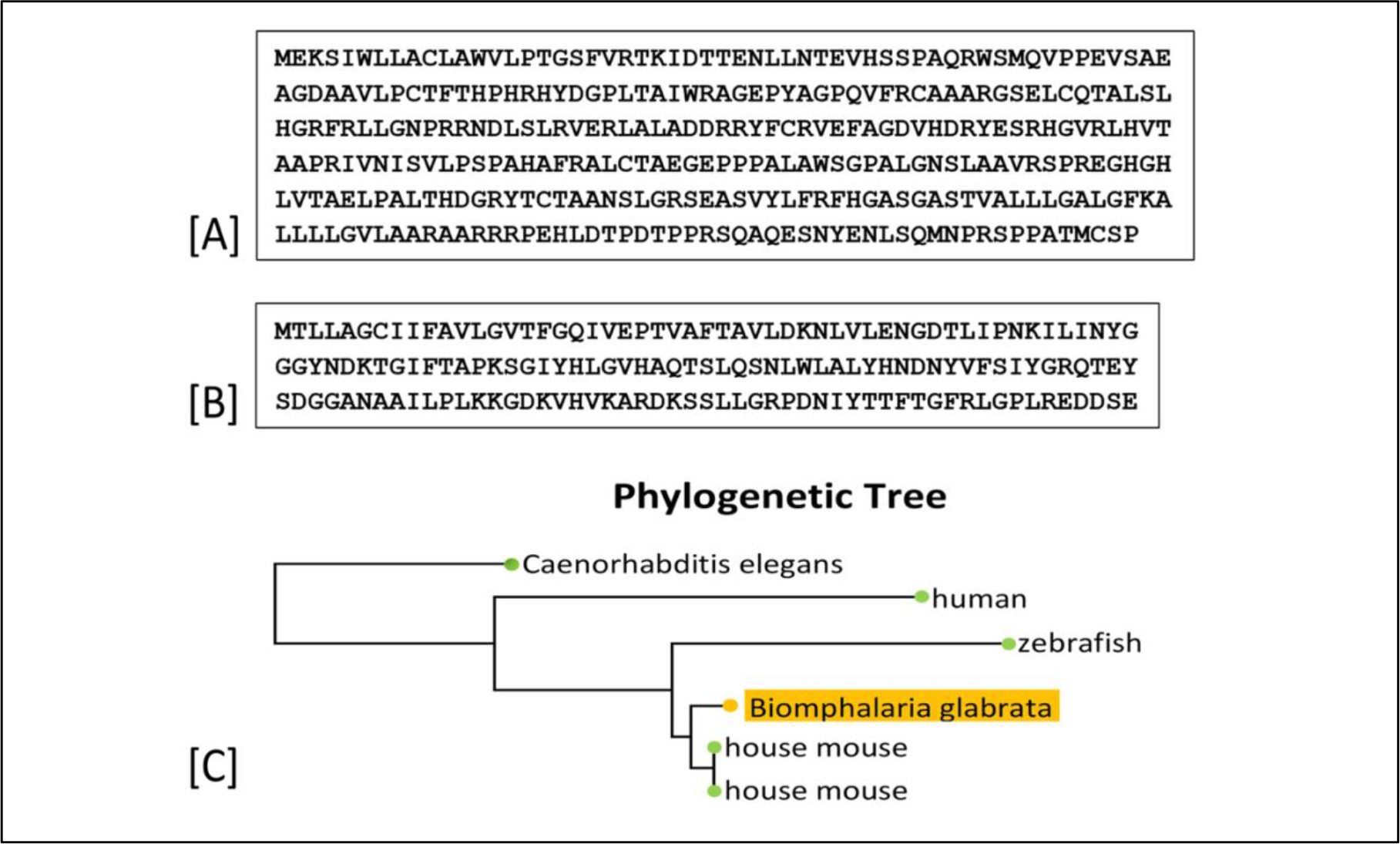
Reference sequences and phylogenetic analysis used to identify *Biomphalaria glabrata* siglec homologs. **(A)** Amino acid sequence of human Siglec-15 (UniProt ID: Q6ZMC9). **(B)** Amino acid sequence of molluscan sialic acid-binding lectin (SABL) from *Cepaea hortensis* (UniProt ID: CAD83837.1). These sequences were used as BLASTp queries to identify *B. glabrata* homologs. **(C)** Phylogenetic tree illustrating the evolutionary relationship between the identified *B. glabrata* siglec homologs and their vertebrate and invertebrate counterparts, showing greater similarity between *B. glabrata* peroxidasin and vertebrate siglecs.

Primers designed from corresponding nucleotides of the proteins listed in Table 1 were utilized for qPCR analysis to determine differences in RNA expression in susceptible BBO2 snails after schistosome exposure for 0, 0.5h,1.0h, 2.0h, 4.0hr, 16.0hr. In Figure 5, results showed by using *B. glabrata* primers for C1q, (homolog of mollusc, *Haliotis discus discus* SALB) early upregulation of this transcript after 1hr exposure to *S. mansoni*, that increased (31-fold upregulation) at 16 hours post-exposure (Figure 5. A). For the *B. glabrata* amino acid sequence showing homology to NR (C1q tumor necrosis factor-related protein 7), results showed (Figure 5. B), a 5-fold upregulation at 30min post-exposure, increasing up to 13-fold within an hour after infection. The transcript showed slight down regulation from 2 to 4 hours, with an overnight increase (10-fold change). For the human siglec15 homolog, peroxidasin, (Figure 5. C) this transcript showed upregulation (1.5-fold) when normalized against the expression of the housekeeping myoglobin encoding transcript for up to 4hr post-parasite exposure in juvenile susceptible BBO2 snails before being downregulated at 16h after infection.

**Figure 4.**
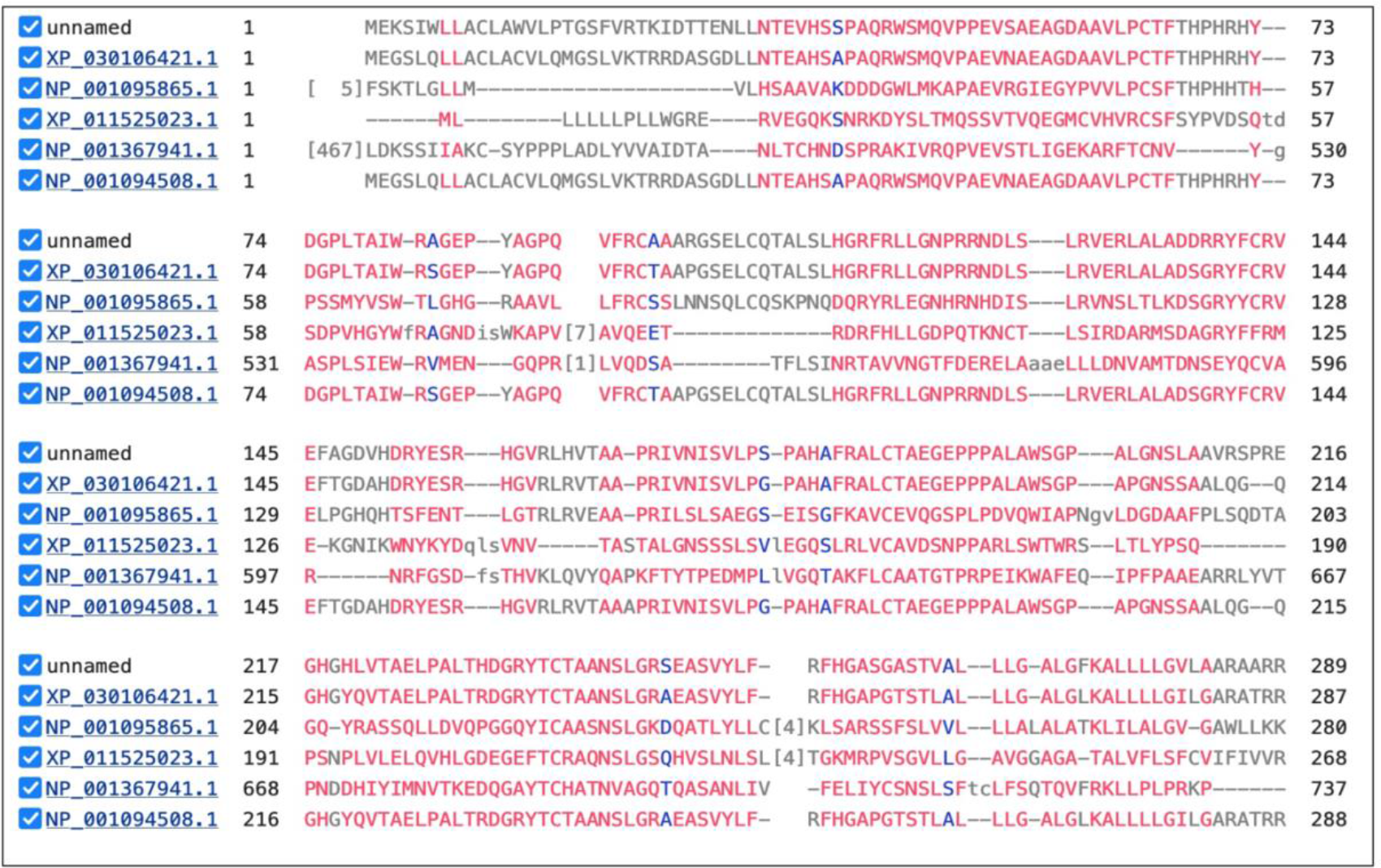
Multiple alignment of B. glabrata siglec with homologs of other organisms. Conserved regions, shown in red, indicate that despite the evolutionary distance between *B. glabrata* peroxidasin and these other siglecs (represented by accession numbers) the relationship between the snail and vertebrate homologs are significant.

**Figure 5.**
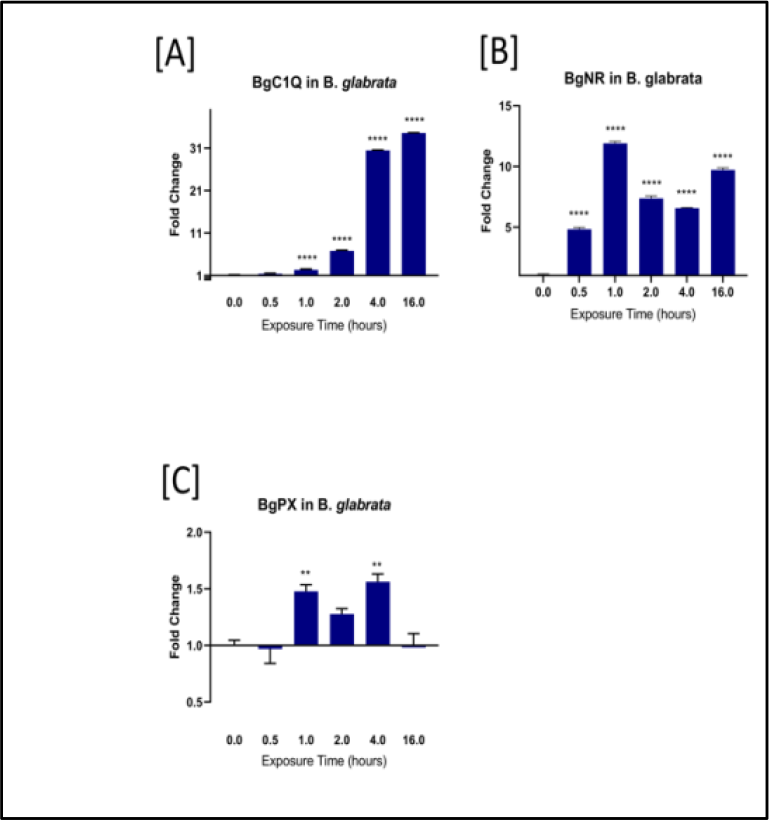
qPCR analysis of RNA from susceptible *B. glabrata* snails exposed to *S. mansoni* miracidia for increasing time-points (0.5 to 16 hours). Histograms show the expression of siglec homolog encoding transcripts in BBO2 snails at specific time points from four biological replicates (the total snails used for four biological replicates n=72). Fold change was determined as described previously by utilizing uniform expression of the reference transcript., ****, p ≤ 0.0001, ***, p ≤ 0.001, **, p ≤ 0.01, *, p ≤ 0.05, and ns, p > 0.05. **(A)** Expression of *Bg*C1q-encoding RNA transcript in susceptible BBO2 snails exposed to *S. mansoni* miracidia; **(B)** Expression of *Bg*NR-encoding RNA transcript in susceptible BBO2 snails exposed to *S. mansoni* miracidia; **(C)** Expression of *Bg*Pxr-encoding RNA transcript in susceptible BBO2 snails exposed to *S. mansoni* miracidia.

### Spatiotemporal regulation of Hsp70 during early *S. mansoni* – *B. glabrata* interaction

Our combined Western blot and immunocytochemistry analyses showed that Hsp70 expression in *Biomphalaria glabrata* upregulates and relocalizes in response to *Schistosoma mansoni* exposure. Quantitative Western blot revealed an up-regulation of cytosolic Hsp70 levels by 1 to 2 hours post-infection. Notably, by the 2-hour time point, an increase in membrane-associated Hsp70 became evident, suggesting a potential relocalization of Hsp70 to the membrane (Figure 6. A and B). Immunocytochemistry of permeabilized *Biomphalaria glabrata* embryonic (*Bge*) cells revealed Hsp70 expression primarily in the cytoplasm of both unexposed and exposed cells, consistent with its known role as a cytosolic chaperone. However, following exposure to *S. mansoni*, a subset of cells revealed enhanced Hsp70 signal near the cell periphery, indicative of partial translocation towards the cell membrane (Figure 6. B and C). This observation aligns with our Western blot data.

**Figure 6.**
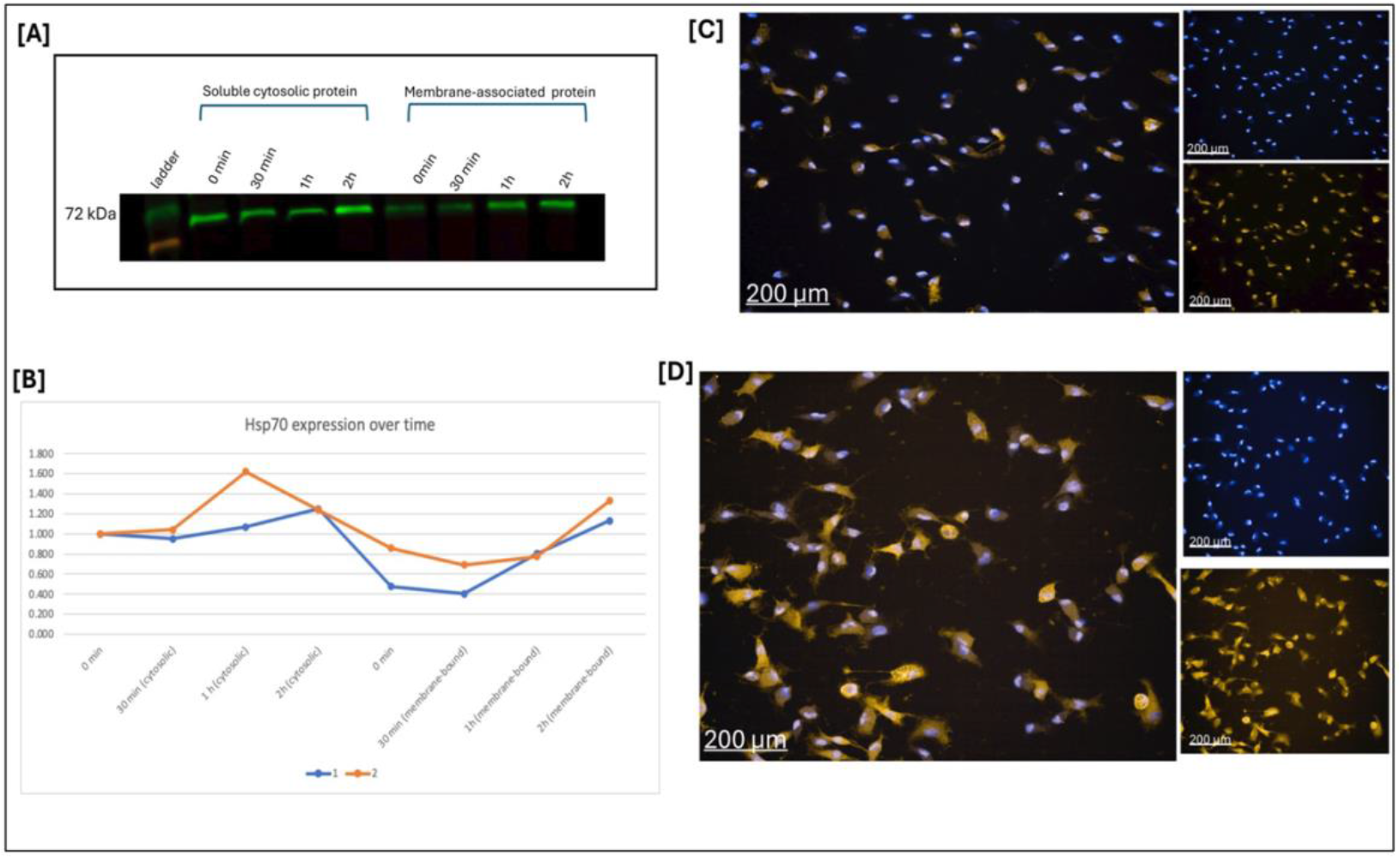
Hsp70 expression dynamics in *B. glabrata* following *S. mansoni* exposure. **(A)** Western blot showing Hsp70 protein levels in cytosolic and membrane-associated protein fractions isolated from juvenile *B. glabrata* snails at 0 min, 30 min, 1 h, and 2 h post-exposure to *S. mansoni* miracidia. Densitometric quantification of Hsp70 band intensities was performed and normalized to total protein. **(B)** Quantification of Hsp70 band intensity from two biological replicates. **(C)** Immunocytochemistry of uninfected *B. glabrata embryonic* (*Bge*) cells showing Hsp70 (orange; Alexa Fluor 555) and nuclei (blue; Hoechst). Merged and individual channels shown. **(D)** *Bge* cells fixed 2 h after exposure to 10 - 15 *S. mansoni* miracidia, showing increased Hsp70 localization at or near the membrane.

Localization of Hsp70 in the cell membrane has previously been reported, but this is the first time that we have shown this is also the case for this stress protein in *B. glabrata*. Notably, surface expression of Hsp70 protein showed a temporal increase for the duration of infection beginning as early as 30 min and increasing to later time points post-exposure (Figure 6. B). Since we have previously shown that the transcript for this stress protein shows an upregulation in exposed susceptible but not resistant snails, the increase in the surface membrane expression of Hsp70 was not surprising. However, the location of the protein in the surface membrane was not expected.

### Computational docking models reveals specific interaction between *BgHsp70* and *BgPrx*

Quality assessments of the homology models indicated favorable stereochemistry. The *BgPrx* model displayed 90.45% of residues in favored Ramachandran regions, a MolProbity score of 1.32, and a GMQE of 0.89. The *BgHsp70* model showed 97% of residues in favored regions, only 0.79% Ramachandran outliers, 0.37% rotamer outliers, a MolProbity score of 0.95, and a GMQE of 0.88. These results support their reliability for downstream docking analysis.

To explore potential interactions between *BgHsp70* and *BgPrx*, we analyzed six docking models that we generated using the PRODIGY web server [28, 29]. This analysis provided binding free energy (ΔG), dissociation constant (Kd), and interfacial contact (IC) data for each complex. Binding free energies ranged from −14.1 to −21.6 kcal/mol, with Models 3 and 4 exhibiting the most favorable interactions (ΔG = −21.6 kcal/mol in both models). Models 3 and 4 also had the lowest predicted Kd values-1.4 × 10^−16^ M and 1.5 × 10^−16^ M, respectively, suggesting *exceptionally* stable complexes. In contrast, the other four models showed weaker interactions, indicating a more transient binding (Table 3).

**Table 3.**
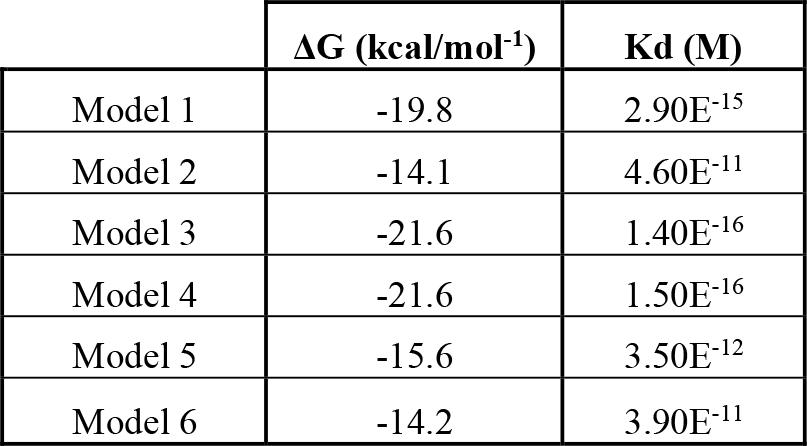
Summary of the binding free energies (ΔG) and dissociation constants (Kd) for each of the six docking models. Models 3 and 4 exhibited the strongest predicted binding affinities, with ΔG values of −21.6 kcal/mol and sub-femtomolar Kd values (∼10^−16^ M), indicating highly stable complexes.

| | $\Delta G$ (kcal/mol<sup>-1</sup>) | $K_d$ (M) |
| --- | --- | --- |
| Model 1 | -19.8 | $2.90E^{-15}$ |
| Model 2 | -14.1 | $4.60E^{-11}$ |
| Model 3 | -21.6 | $1.40E^{-16}$ |
| Model 4 | -21.6 | $1.50E^{-16}$ |
| Model 5 | -15.6 | $3.50E^{-12}$ |
| Model 6 | -14.2 | $3.90E^{-11}$ |

Given its favorable ΔG and Kd, we selected Model 3 for detailed analysis using Mol*, a web-based visualization tool [30]. A prominent hydrogen bond was observed between LYS47 on *BgPrx* and GLU479 on *BgHsp70*, forming a classic salt bridge interaction (Figure 7. D). This specific pairing aligns well with previous findings [27], which showed that positively charged residues such as Arg119 and Lys134 within the V-set domain of human Siglec-14 are essential for binding Hsp70. While that study did not identify the corresponding contact residues on Hsp70, our model reveals a structurally plausible polar–polar interface. Notably, LYS47 lies at the amino terminus of a β-strand that forms part of a V-set domain (Figure 7. C).

**Figure 7.**
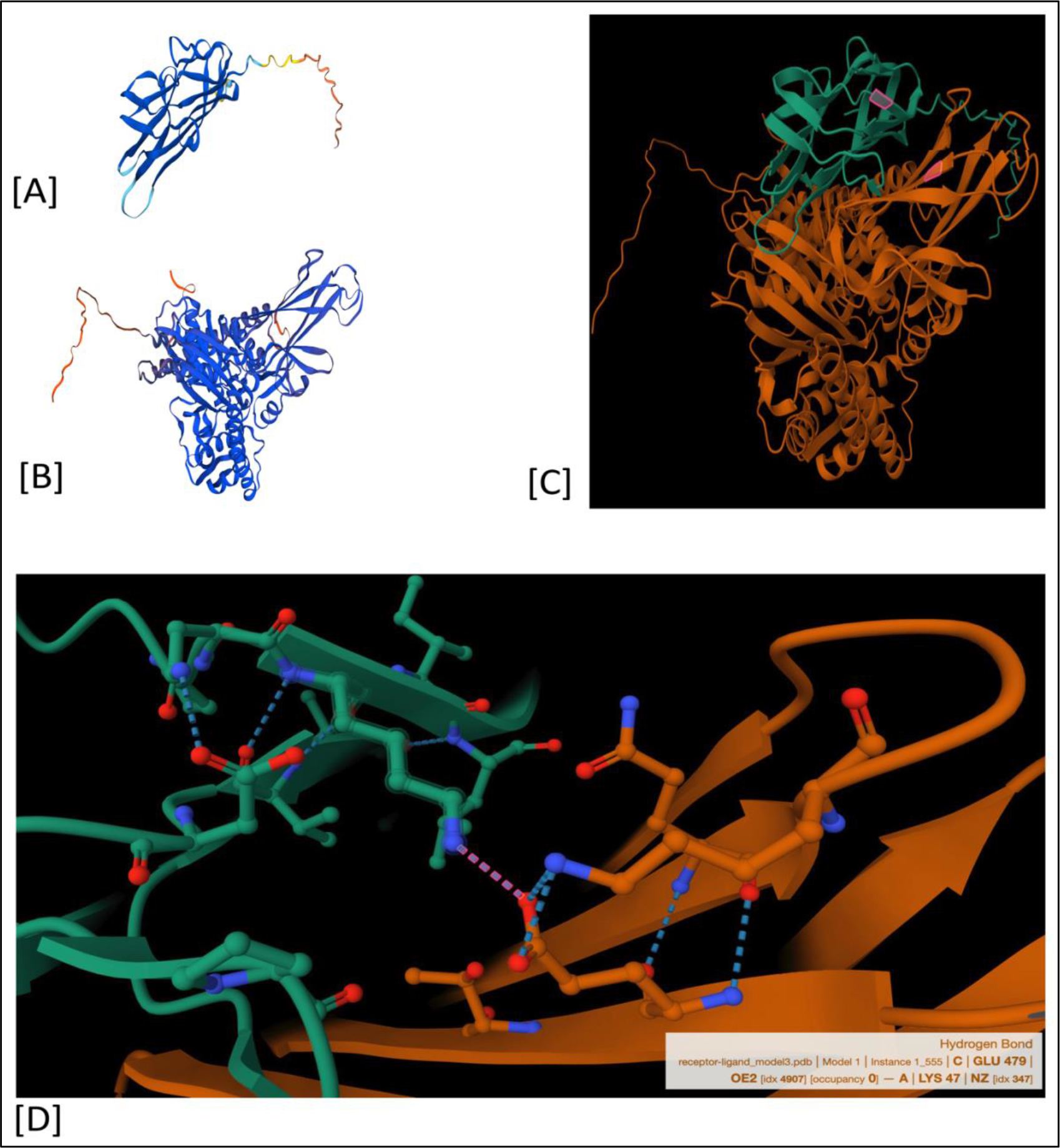
Homology Modelling and Docking Reveal Salt Bridge Between *BgHsp70* and *BgPrx*. **(A)** Homology model of *Biomphalaria glabrata* peroxidasin-like protein (*BgPrx*), generated using SWISS-MODEL and displayed in cartoon representation with N- to C-terminal gradient coloring. **(B)** Homology model of *B. glabrata* heat shock protein 70(*BgHsp70)*. **(C)** Docked complex of *BgHsp70* (orange) and *BgPrx* (green) illustrating their predicted interaction interface. The pink highlight marks the region of interest showing a potential salt bridge between GLU 479 of *BgHsp70* and LYS 47 of *BgPrx*. **(D)** Close-up view of the predicted salt bridge at the binding interface. Hydrogen bonding is shown as pink dashed lines, with a key interaction occurring between the OE2 atom of GLU 479 (*BgHsp70*) and the NZ atom of LYS 47 *(BgPrx)*, indicating a salt bridge that may contribute to binding affinity and complex stability.

These data show that with limited information, existing sequences in public domains can readily be accessed to search for unknown molecular determinants that play a role in modulating innate immunity in understudied organisms, such as *B. glabrata*. With genomic sequences from many organisms now becoming available at a pace that was unimagined a decade ago, we are in a new era where we can identify functional transcripts starting with very little information as long there are publicly funded and well-curated databases at our disposal. With this in-silico approach, we have shown here that we were able to identify potentially relevant immune response transcripts that are related to human and molluscan siglecs in *B. glabrata*. Using previous information that showed Hsp70 plays a role in innate defence and information that this stress protein also binds to human siglecs [18], it was a logical step to search for siglec homologs in *B. glabrata*. It is clear from our study, that *B. glabrata* Hsp70 protein shows a membrane location that increases with time post-infection. We are using pull-down co-immunoprecipitation assays, to identify client proteins, followed by a proteomic approach to reveal the identity of proteins that bind to Hsp70 in early exposed susceptible *B. glabrata* juvenile snails. *B. glabrata* C1q has previously been characterized and so have the family of over 30 NR genes ([22]), however, *B. glabrata* peroxidasin and its significant homology to human siglec15 is shown here for the first time. Human siglec 15 is expressed on macrophages and is also important in cancer, especially pancreatic cancer. It has recently been shown that human siglec 15 is a cancer immunosuppressor that is being targeted for cancer immunotherapy [23].

In future studies, we plan to examine, by qPCR, the occurrence of siglec homologs identified in this study in the other medically important snails that also transmit schistosomiasis, namely snails of *Bulinus* and *Oncomelania* genera that serve as hosts for *Schistosoma haematobium* and *Schistosoma japonicum*, respectively. *S. haematobium*, causes urinary schistosomiasis and is classified by International Agency for Research on Cancer (IARC) as a grade 1 infectious agent that causes bladder, cervical cancer and both male and female infertility [24, 25]. *S. japonicum* causes intestinal schistosomiasis and colorectal cancer [25]. Using antibodies against the siglec homologs identified in this study, we will examine in more detail, from spatial and temporal studies, the interaction of these proteins with Hsp70. We hope that these studies will provide a better understanding of how stress proteins and innate defence are linked in the anti-schistosome response in the *B. glabrata* host-schistosome relationship.

## Acknowledgement

We thank Adeola Fagunloye for designing the *Schistosoma mansoni* lifecycle figure and Lidia Campos-Zurita for the illustration showing the association of a siglec ligand and the membrane-bound Hsp70. This work was supported in part by the Clement B.T. Knight Cancer Foundation and a grant to Drs. Freddie Dixon and Carolyn Cousin from the National Science Foundation Grant (Award No. 1622811). The BBO2 susceptible snails were provided by the Schistosomiasis Resource Center of the Biomedical Research Institute (Rockville, MD) through NIH-NIAID Contract HHSN272201700014I.

## Notes

### Competing Interest Statement

The authors have declared no competing interest.

### Summary of Updates

This revised version includes a correction to the author name (Matty Knight, previously displayed with a middle initial) and a minor edit to the abstract to incorporate the role of glycan mimicry in immune evasion. No additional changes were made to the data, figures, or main body of the manuscript.

## REFERENCES

1 Colley DG, Bustinduy AL, Secor WE, King CH. Human schistosomiasis. Lancet. 2014;383(9936):2253–64. Epub 2014/04/05. doi: 10.1016/S0140-6736(13)61949-2. PubMed PMID: 24698483; PubMed Central PMCID: PMCPMC4672382.

2 LoVerde PT. Schistosomiasis. Adv Exp Med Biol. 2019;1154:45–70. doi: 10.1007/978-3-030-18616-6_3. PubMed PMID: 31297759.

3 Ogongo P, Nyakundi RK, Chege GK, Ochola L. The Road to Elimination: Current State of Schistosomiasis Research and Progress Towards the End Game. Front Immunol. 2022;13:846108. Epub 20220503. doi: 10.3389/fimmu.2022.846108. PubMed PMID: 35592327; PubMed Central PMCID: PMCPMC9112563.

4 Hambrook JR, Hanington PC. Immune Evasion Strategies of Schistosomes. Front Immunol. 2020;11:624178. Epub 20210204. doi: 10.3389/fimmu.2020.624178. PubMed PMID: 33613562; PubMed Central PMCID: PMCPMC7889519.

5 Ittiprasert W, Nene R, Miller A, Raghavan N, Lewis F, Hodgson J, et al. Schistosoma mansoni infection of juvenile Biomphalaria glabrata induces a differential stress response between resistant and susceptible snails. Exp Parasitol. 2009;123(3):203–11. Epub 2009/08/08. doi: 10.1016/j.exppara.2009.07.015. PubMed PMID: 19660454; PubMed Central PMCID: PMCPMC2760455.

6 Ittiprasert W, Knight M. Reversing the resistance phenotype of the Biomphalaria glabrata snail host Schistosoma mansoni infection by temperature modulation. PLoS Pathog. 2012;8(4):e1002677. Epub 20120426. doi: 10.1371/journal.ppat.1002677. PubMed PMID: 22577362; PubMed Central PMCID: PMCPMC3343117.

7 Knight M, Elhelu O, Smith M, Haugen B, Miller A, Raghavan N, et al. Susceptibility of Snails to Infection with Schistosomes is influenced by Temperature and Expression of Heat Shock Proteins. Epidemiology (Sunnyvale). 2015;5(2). Epub 20150621. doi: 10.4172/2161-1165.1000189. PubMed PMID: 26504668; PubMed Central PMCID: PMCPMC4618387.

8 Smith M, Yadav S, Fagunloye OG, Pels NA, Horton DA, Alsultan N, et al. PIWI silencing mechanism involving the retrotransposon nimbus orchestrates resistance to infection with Schistosoma mansoni in the snail vector, Biomphalaria glabrata. PLoS Negl Trop Dis. 2021;15(9):e0009094. Epub 20210908. doi: 10.1371/journal.pntd.0009094. PubMed PMID: 34495959; PubMed Central PMCID: PMCPMC8462715.

9 Dinguirard N, Cavalcanti MGS, Wu XJ, Bickham-Wright U, Sabat G, Yoshino TP. Proteomic Analysis of Biomphalaria glabrata Hemocytes During in vitro Encapsulation of Schistosoma mansoni Sporocysts. Front Immunol. 2018;9:2773. Epub 2018/12/18. doi: 10.3389/fimmu.2018.02773. PubMed PMID: 30555466; PubMed Central PMCID: PMCPMC6281880.

10 Yoshino TP, Boyle JP, Humphries JE. Receptor-ligand interactions and cellular signalling at the host-parasite interface. Parasitology. 2001;123 Suppl:S143-57. PubMed PMID: 11769279.

11 Seppala O, Cetin C, Cereghetti T, Feulner PGD, Adema CM. Examining adaptive evolution of immune activity: opportunities provided by gastropods in the age of ‘omics’. Philos Trans R Soc Lond B Biol Sci. 2021;376(1825):20200158. Epub 20210405. doi: 10.1098/rstb.2020.0158. PubMed PMID: 33813886; PubMed Central PMCID: PMCPMC8059600.

12 Lockyer AE, Spinks JN, Walker AJ, Kane RA, Noble LR, Rollinson D, et al. Biomphalaria glabrata transcriptome: identification of cell-signalling, transcriptional control and immune-related genes from open reading frame expressed sequence tags (ORESTES). Dev Comp Immunol. 2007;31(8):763-82. PubMed PMID: 17208299.

13 Raghavan N, Miller AN, Gardner M, FitzGerald PC, Kerlavage AR, Johnston DA, et al. Comparative gene analysis of Biomphalaria glabrata hemocytes pre-and post-exposure to miracidia of Schistosoma mansoni. Mol Biochem Parasitol. 2003;126(2):181-91. PubMed PMID: 12615317.

14 Tennessen JA, Bollmann SR, Peremyslova E, Kronmiller BA, Sergi C, Hamali B, et al. Clusters of polymorphic transmembrane genes control resistance to schistosomes in snail vectors. Elife. 2020;9. Epub 20200826. doi: 10.7554/eLife.59395. PubMed PMID: 32845238; PubMed Central PMCID: PMCPMC7494358.

15 Adema Cmal, E.S. Specificity and immunobiology of larval digenean snail associations. In: Fried, B Graczyk, T.K. (Eds.) Advances in Trematode Biology. CRC Press, Boca Raton, FL. 1997:229–63.

16 Zhang SM, Zeng Y, Loker ES. Characterization of immune genes from the schistosome host snail Biomphalaria glabrata that encode peptidoglycan recognition proteins and gram-negative bacteria binding protein. Immunogenetics. 2007;59(11):883-98. PubMed PMID: 17805526.

17 Adema CM, Hillier LW, Jones CS, Loker ES, Knight M, Minx P, et al. Corrigendum: Whole genome analysis of a schistosomiasis-transmitting freshwater snail. Nat Commun. 2017;8:16153. Epub 2017/08/24. doi: 10.1038/ncomms16153. PubMed PMID: 28832025; PubMed Central PMCID: PMCPMC5569240.

18 Fong JJ, Sreedhara K, Deng L, Varki NM, Angata T, Liu Q, et al. Immunomodulatory activity of extracellular Hsp70 mediated via paired receptors Siglec-5 and Siglec-14. EMBO J. 2015;34(22):2775–88. Epub 20151012. doi: 10.15252/embj.201591407. PubMed PMID: 26459514; PubMed Central PMCID: PMCPMC4682649.

19 Livak KJ, Schmittgen TD. Analysis of relative gene expression data using real-time quantitative PCR and the 2(-Delta Delta C(T)) Method. Methods. 2001;25(4):402–8. doi: 10.1006/meth.2001.1262. PubMed PMID: 11846609.

20 Knight M, Raghavan N, Goodall C, Cousin C, Ittiprasert W, Sayed A, et al. Biomphalaria glabrata peroxiredoxin: effect of schistosoma mansoni infection on differential gene regulation. Mol Biochem Parasitol. 2009;167(1):20–31. Epub 2009/05/15. doi: 10.1016/j.molbiopara.2009.04.002. PubMed PMID: 19439374; PubMed Central PMCID: PMCPMC2705950.

21 Myers J, Ittiprasert W, Raghavan N, Miller A, Knight M. Differences in cysteine protease activity in Schistosoma mansoni-resistant and -susceptible Biomphalaria glabrata and characterization of the hepatopancreas cathepsin B Full-length cDNA. J Parasitol. 2008;94(3):659-68. PubMed PMID: 18605796.

22 Kaur S, Jobling S, Jones CS, Noble LR, Routledge EJ, Lockyer AE. The nuclear receptors of Biomphalaria glabrata and Lottia gigantea: implications for developing new model organisms. PLoS One. 2015;10(4):e0121259. Epub 20150407. doi: 10.1371/journal.pone.0121259. PubMed PMID: 25849443; PubMed Central PMCID: PMCPMC4388693.

23 Sun J, Lu Q, Sanmamed MF, Wang J. Siglec-15 as an Emerging Target for Next-generation Cancer Immunotherapy. Clin Cancer Res. 2021;27(3):680–8. Epub 20200921. doi: 10.1158/1078-0432.CCR-19-2925. PubMed PMID: 32958700; PubMed Central PMCID: PMCPMC9942711.

24 Jenkins-Holick DS, Kaul TL. Schistosomiasis. Urol Nurs. 2013;33(4):163-70. PubMed PMID: 24079113.

25 Ishii A, Matsuoka H, Aji T, Ohta N, Arimoto S, Wataya Y, et al. Parasite infection and cancer: with special emphasis on Schistosoma japonicum infections (Trematoda). A review. Mutat Res. 1994;305(2):273–81. doi: 10.1016/0027-5107(94)90247-x. PubMed PMID: 7510038.

26 Singh, A., Copeland, M.M., Kundrotas, P.J., Vakser, I.A., 2024, GRAMM Web Server for Protein Docking, Methods in Mol. Biol., 2714:101–112.

27 Fong JJ, Nguyen D, Dharmarajan V, Agrawal S, Tran D, Randhawa V, et al. Siglec-5 and Siglec-14 are receptors for the endoplasmic reticulum chaperone Hsp70. EMBO J. 2015;34(24):2775–88. doi:10.15252/embj.201592394. PMID: 26424579; PMCID: PMC4682649.

28 Vangone A. and Bonvin A.M.J.J. “Contact-based prediction of binding affinity in protein-protein complexes”, eLife, 4, e07454 (2015).

29 Xue L., Rodrigues J., Kastritis P., Bonvin A.M.J.J.*, Vangone A.*, “PRODIGY: a web-server for predicting the binding affinity in protein-protein complexes”, Bioinformatics, doi:10.1093/bioinformatics/btw514 (2016).

30 Sehnal D, Bittrich S, Deshpande M, Svobodová R, Berka K, Bazgier V, et al. Mol*: Towards a common library and tools for web molecular graphics. MolVA’18: Workshop on Molecular Graphics and Visual Analysis of Molecular Data; 2018. p. 29–33. doi:10.2312/molva.20181103.

31 Waterhouse A, Bertoni M, Bienert S, Studer G, Tauriello G, Gumienny R, et al. SWISS-MODEL: homology modelling of protein structures and complexes. Nucleic Acids Res. 2018;46(W1):W296–303. doi:10.1093/nar/gky427. PMID: 29788355; PMCID: PMC6030848.

